# Looking for unique signals of human expansion from Africa: Beyond diversity

**DOI:** 10.1101/2024.08.13.607798

**Authors:** Zarus Cenac

## Abstract

There are known to be different views on which portion of Africa modern humans globally spread from. Biological diversities have been employed to estimate the origin of our global expansion. These diversities vary with geographical distance from Africa, thereby expressing the signal of the expansion. In preprints, the signal supposedly appeared beyond diversity – in cranial sexual size dimorphism and a cranial shape distance-based measure. Compared to when diversity is used alone, the addition to analysis of variables which are beyond diversity could improve recovery of the signal, therefore improving origin identification. I explored this through cranial and genetic measures which had been calculated in prior studies. Various analyses were used, e.g., ridge regression and Mantel tests. Amongst cranial variables (shape diversity, sexual size dimorphism, and a shape distance-based measure), only dimorphism had a unique portion of the expansion signal. In comparison to when diversity was utilised alone, the additional use of dimorphism and the distance-based measure did not substantially impact signal recovery. However, their addition possibly improved origin identification, reducing by 46% the size of the geographical area which may have the origin. This smaller area approximately matched southern Africa, however, it was not only in the south. It was questionable if the signal was present in a genetic distance-based measure, which called into question whether the expansion signal is truly present in the cranial shape distance-based measure. Analysis suggested that the apparent presence of the signal in distance-based measures is affected by the representation of Oceanian populations. This study supports cranial sexual size dimorphism being a helpful indicator of the expansion whilst calling into question whether biological distance-based measures are indicators. Clarity remains missing on which African region was the origin.

## Introduction

Modern humans originated in Africa, whether in a single region or from multiple regions of the continent (e.g., Henn et al., 2018). They spread out, across the continent and beyond, dispersing across the globe through population bottlenecks (Henn et al., 2012; Ramachandran et al., 2005), with genetic drift consequently increasing along the way (Ramachandran et al., 2005). The effect of these bottlenecks seems to be apparent in genetic diversity, which decreases as distance from Africa increases (Ramachandran et al., 2005); the signal of the expansion in genetics is reflected in the cranium (Betti et al., 2009), with cranial diversity also lessening as distance from the continent increases (Manica et al., 2007; von Cramon Taubadel & Lycett, 2008) (see Figure 1).

**Figure 1.**
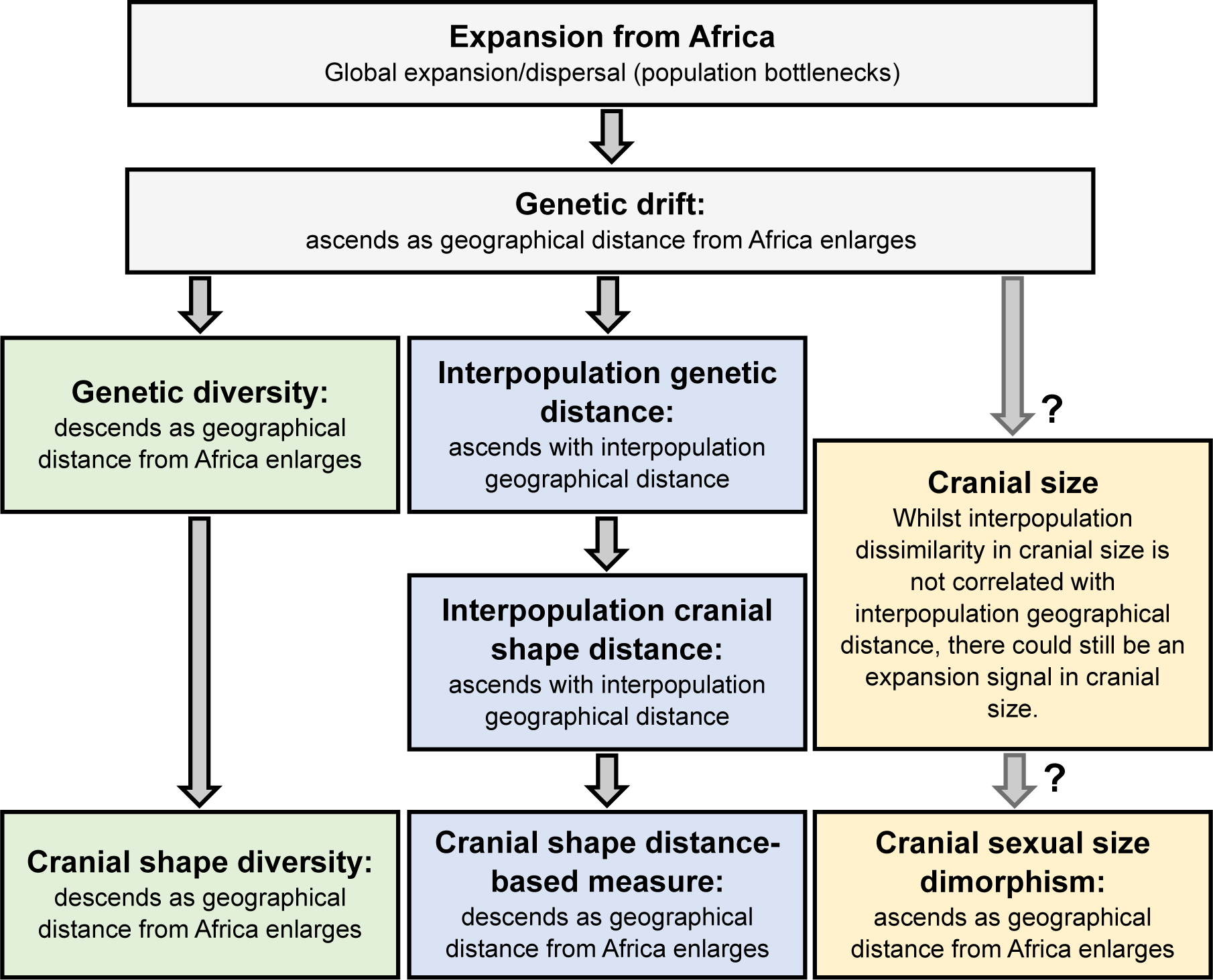
Effects of Global Expansion on Genetics and Crania. *Notes*. Figure 1 is based on research and discourse (Betti et al., 2009, 2010; Cenac, 2022a, 2022b, 2023b, 2024; Harvati & Weaver, 2006; Hubbe et al., 2009; Ponce de León et al., 2018; Ramachandran et al., 2005; von Cramon-Taubadel & Lycett, 2008). Expansion from an origin suggests why i) the genetic diversity of populations goes downward the farther geographically populations are from Africa, and ii) populations are less genetically alike as the geographical distance between these populations greatens (Ramachandran et al., 2005). The expansion signal in genetics is evident in the cranium (Betti et al., 2009), with cranial form/shape diversity reducing as distance from Africa increases (Betti et al., 2009; von Cramon-Taubadel & Lycett, 2008); the signal in cranial form diversity is driven by the signal in cranial shape diversity – cranial size diversity does not exhibit the signal (Cenac, 2022a). The farther the geographical distance between populations, the more biologically dissimilar these populations are not only when measured in genetics (e.g., Ramachandran et al., 2005), but also in terms of cranial shape (Hubbe et al., 2010), but not in cranial size (Harvati & Weaver, 2006); the relationship regarding cranial shape difference (plus the expansion having followed various routes) may lead to the expansion signal appearing supposedly in a cranial shape distance-based measure (for the male cranium) (Cenac, 2023b). It is not apparent why cranial sexual size dimorphism has a positive correlation with geographical distance from Africa, with it not being clear if the expansion signal is of greater prominence in the cranial size of males than females (Cenac, 2024).

### The global expansion: Its signal and origin

The expansion signal seems to be stronger in microsatellite diversity than in cranial diversity (Roseman & Weaver, 2007). Indeed, distance from Africa explains about 85% of the variance in autosomal microsatellite heterozygosity (Prugnolle et al., 2005; Ramachandran et al., 2005), but only about 20-30% of the variance in cranial form and shape diversities (Betti et al., 2009; von Cramon-Taubadel & Lycett, 2008). It does appear that the genetic-derived signal (and therefore expansion signal) can be recovered better across particular measurements (10 dimensions) from the front of the cranium rather than from the cranium overall, with distance from Africa explaining 50% of the variance of these particular measurements (Betti et al., 2009). However, even with these traits, the cranium stills falls short when compared to genetics (Betti et al., 2009).

Whilst Africa seems to be where the expansion originated (Ramachandran et al., 2005), there are different positions on which region of the continent was the origin (e.g., Ray et al., 2005: Tishkoff et al., 2009), with the south and the east seeming to be regarded as the frontrunners (Henn et al., 2018). Genetic and cranial diversities have been used to identify where the expansion originated (e.g., Manica et al., 2007). There are a number of ways of identifying the origin (e.g., Betti et al., 2009; Ramachandran et al., 2005; Ray et al., 2005; Tishkoff et al., 2009). For instance, a method identifies a range of locations amongst which lies the origin (Manica et al., 2007; as an example, see Figure 2A).

**Figure 2.**
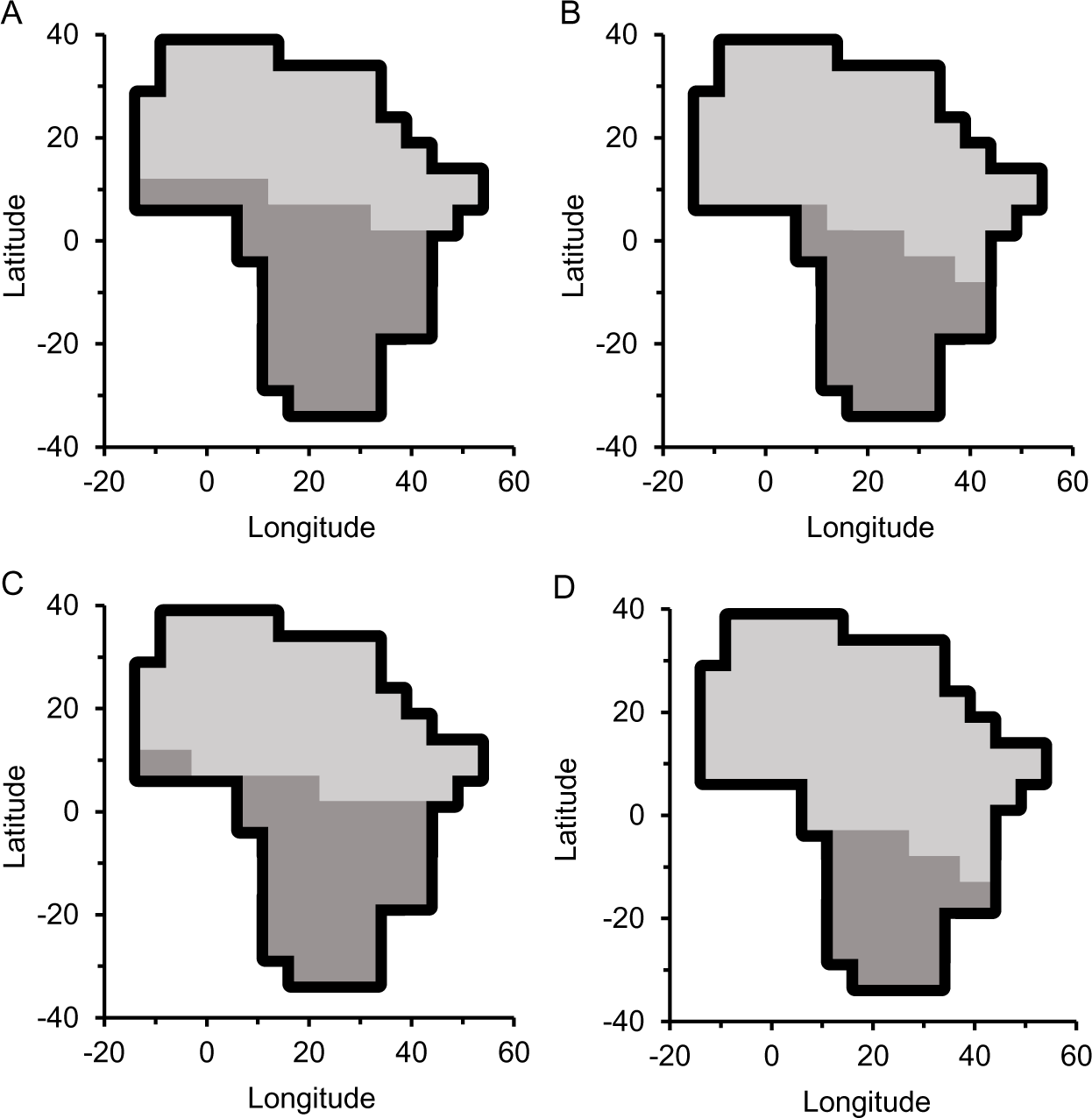
Cranium: Identification of Where the Expansion Originated. *Note*. Figure 2A is with respect to cranial shape diversity, Figure 2B used the diversity alongside cranial sexual size dimorphism, Figure 2C used diversity with the cranial shape distance-based measure, and Figure 2D used cranial shape diversity, cranial sexual size dimorphism, and the cranial shape distance-based measure. The darker grey represents the area in which the origin is estimated to be, unlike the lighter grey area.

Recovering a stronger expansion signal may allow for a more precise identification of the origin. Signal strength could be improved by adding a variable (or variables) to an analysis (e.g., Betti et al., 2009). Ideally, the added variable would have a unique portion of the expansion signal (compared to any variable already in the analysis) (e.g., Betti et al., 2009). Perhaps then, signal recovery would be served well by using variables in an analysis which are as different as possible from each other – perhaps diversity could be used alongside variables which are beyond diversity (see Figure 1).

### Beyond diversity: Cranial sexual size dimorphism and interpopulation biological distance

Indeed, selecting a subset of measurements (Betti et al., 2009) may not be the only way of recovering a stronger expansion signal in the cranium. Alongside cranial diversity, other cranially-derived traits which have the expansion signal (Cenac, 2022b, 2023b) could be considered. Interestingly, the cranium appears to not only have the expansion signal in diversity (Betti et al., 2009; Manica et al., 2007; von Cramon-Taubadel & Lycett, 2008), but preprints suggest that the signal is present in cranial sexual size dimorphism (Cenac, 2022b) and, for male crania, a measure based on interpopulation cranial shape distances – a **cranial shape distance-based measure** (Cenac, 2023b) (see *Glossary*; Figure 1). Distance from Africa explains about 45% and 20% of the respective variance in cranial sexual size dimorphism and the cranial shape distance-based measure (Cenac, 2022b, 2023b).

Linear regression could be used to see if these two cranial measures have distinct parts of the expansion signal (amongst cranial measures). If one or both has a distinct portion of the signal, then the use of three cranially-derived measures (diversity, dimorphism, and shape distance-based) could lead to a stronger recovery of the expansion signal than cranial diversity does alone. Linear regression can be used to infer whether a stronger signal is recovered. If there is a stronger signal, we may expect that these three variables may collectively lead to a smaller (and ideally more precise) geographical area being identified as the area in which the origin resides (compared to diversity alone). A similar analysis could occur with genetic data, i.e., finding whether a **genetic distance-based measure** (Cenac, 2023b) (see *Glossary*) improves origin identification compared to genetic diversity alone.

### Origin identification across variables

Before, in a preprint, I looked into identifying the origin of the expansion by averaging an estimate from various genetic and cranial variables (Cenac, 2023b).^1^ This averaging resulted in something like a heatmap of Africa, ranking how good a fit 99 locations across the continent are at being the origin (with locations ranked 1 to 99), and this averaging indicated a southern African origin (Cenac, 2023b). By using rankings, this method does not carve out a distinctive geographical area of Africa as the area which has the origin. Such a distinctive area has been found through individual genetic and cranial variables, each on their own (e.g., Betti et al., 2009; Manica et al., 2007; see Figure 2A for an example); a distinctive area was not known to have been carved out *across* variables which each indicate the expansion (i.e., considering *different variables*^2^ simultaneously).

### Present study

Because of the of bottlenecks in the expansion from Africa (Ramachandran et al., 2005), and the relationships between geographical distance from Africa and cranial/genetic variables (Cenac, 2022b, 2023b; Prugnolle et al., 2005; von Cramon-Taubadel & Lycett, 2008), geographical distance from the origin of the expansion was used as a proxy for the *true* signal of the expansion.

#### Main Analysis 1

This study tested whether cranial sexual size dimorphism and the cranial shape distance-based measure i) have unique portions of the expansion signal amongst cranial measures, and ii) if they lead to a better recovery of the expansion signal in the cranium compared to when only diversity is considered. It was then seen, descriptively, if their inclusion reduces the size of the geographical area in which the origin is identified. This involved estimating, across cranial variables, a distinct geographical area regarding the origin of the expansion.

#### Main Analysis 2

The expansion signal is stronger in genetic diversity than in cranial diversity (Roseman & Weaver, 2007). Therefore, even if cranial sexual size dimorphism or the cranial shape distance-based measure have a unique part of the signal amongst cranial measures, they could still be redundant when compared to autosomal diversity. Given that the genetic-derived signal is thought to be represented in cranial diversity (Betti et al., 2009), cranial diversity would not be expected to have a segment of the expansion signal which is distinct to the signal in genetic diversity. Therefore, it was seen if the three cranial measures (sexual size dimorphism, the shape distance-based measure, and shape diversity) have portions of the expansion signal which are distinct to not only each other but also distinct to autosomal microsatellite heterozygosity.

#### Main Analysis 3

An issue with using the cranial shape distance-based measure to identify the origin is nonindependence – this could be allayed by seeing if a genetic distance-based measure also suggests the origin of the expansion (Cenac, 2023b). The present study explored this.

#### Main Analysis 4

Additionally, this study tested i) if the genetic distance-based measure has some of the expansion signal which is not present in autosomal diversity, and ii) whether the expansion signal in genetics could be recovered better (vs. autosomal diversity alone) by using the genetic distance-based measure alongside genetic diversity, thereby potentially improving the identification of the origin. In an additional (unplanned) analysis, it was seen if iii) compared to a number of genetic diversities (autosomal, mitochondrial, and X-chromosomal), whether a stronger expansion signal is recovered by additionally considering the genetic distance-based measure.

## Method

The present study used data from previous research. Calculated from the Howells cranial dataset (Howells, 1973, 1989, 1995, 1996; http://web.utk.edu/~auerbach/HOWL.htm), the present study used i) the within-population shape diversity of males in 26 populations (Cenac, 2023b) and ii) the cranial sexual size dimorphism of 26 populations (Cenac, 2022a). Also calculated from the Howells dataset, this study utilised, iii) the cranial shape distance-based measure (Spearman correlation coefficients) (Cenac, 2023b), and iv) cranial shape measurements (males from 21 populations) calculated from the Howells data (Cenac 2022a, 2022b). As for genetic data, this study used the autosomal microsatellite heterozygosity and X-chromosomal microsatellite heterozygosity of 50 HGDP-CEPH populations (Balloux et al., 2009), and also mitochondrial diversity (from pairwise differences – the mean amount of differences)^3^ from 50 HGDP-CEPH populations (Lippold et al., 2014). Also used were genetic distances between pairs of populations calculated for 49 populations in the HGDP-CEPH data, and another population (Nasioi, who correspond to Melanesians in the HGDP-CEPH data) from autosomal microsatellites (Friedlaender et al., 2008). These distances were used to make a genetic equivalent of the cranial shape distance-based measure, and this genetic distance-based measure was employed in analyses akin to how Cenac (2023b) used the cranial shape distance-based measure. Analyses used geographical distances which included distances from locations that were used as expansion origins, with these origins being 99 ones in Africa alone (from Betti et al., 2013) and 32 which are across the globe (from von Cramon-Taubadel & Lycett, 2008) (Cenac, 2022a, 2022b). Analyses were undertaken in R Version 4.0.5 (R Core Team, 2021) and Microsoft Excel. Figures 2, 6, and 7 (graphs showing a map of Africa) are based, in design, on Cenac (2022), who utilised coordinates shown in a figure in Betti et al. (2013). Figure 2A is like Figure 2E of Cenac (2022b), who also used cranial shape diversity stemming from the Howells data, however, whilst Cenac (2022b) used 28 populations, Figure 2A only featured 24 populations. Figure 6A is similar to Figure 1A of Cenac (2023a) – both used (from Pemberton et al., 2013) the heterozygosity of autosomal microsatellites from populations in the HGDP-CEPH dataset, but, whilst Figure 6A used 49 populations, Figure 1A of Cenac (2023a) used 50 populations. In multiple linear regressions, the presence of multicollinearity was assessed using the car package (Fox & Weisberg, 2019) in R, through which variation inflation factors were calculated. A variance inflation factor (VIF) represents the relationship between a variable and other variables, and a value in excess of 5–10 VIFs portrays the presence of multicollinearity (Kim, 2019).

### Main analyses

*Main Analysis 1* was with respect to predicting distance from Africa from three variables (cranial shape diversity, cranial sexual size dimorphism, and the cranial shape distance-based measure). To perform this analysis, linear regression was used on 26 populations.

*Main Analysis 2* was about the prediction of distance from Africa from four variables. This time autosomal microsatellite heterozygosity was used (Balloux et al., 2009, i.e., originating from HGDP-CEPH) with cranial shape diversity, cranial sexual size dimorphism, and the cranial shape distance-based measure (Cenac, 2022a, 2023b, i.e., originating from the Howells data). This linear regression was performed on distance matrices using MRM in the ecodist package of Goslee and Urban (2007) in R. To perform the analysis, HGDP-CEPH and Howells populations were matched following Roseman (2004). However, there were some differences from Roseman (2004). Firstly, Roseman (2004) used Anyang, whereas the present analysis did not. This was because the Howells (e.g., 1996) Anyang data only features males. Roseman (2004) combined together Melanesians and Papuans regarding HGDP-CEPH data; the present study did not use the Papuan HGDP-CEPH population in the event of doing additional analysis with the genetic distance-based measure (Friedlaender et al., 2008, did not include the Papuan HGDP-CEPH population). Literature was used (Balloux et al., 2009; Hayashi & Pietrusewsky, 2023; Roseman, 2004; von Cramon-Taubadel & Lycett, 2008) to check that matching between HGDP-CEPH and Howells datasets from Roseman (2004) was followed (aside from the Anyang population in the Howells data and the Papuan population in the HGDP-CEPH data). All in all, whilst matching resulted in 10 geographical groupings (each consisting of one Howells population matched to one or more HGDP-CEPH population) in Roseman (2004), matching in the present study resulted in nine geographical groupings. Main Analysis 2 was supplemented (post-hoc) with a further linear regression on distance matrices using ecodist, and three Mantel correlation tests, with each Mantel test being supplemented by a further Mantel correlation test. These six Mantel correlation tests used the Mantel function in the ecodist package. For Mantel tests, geographical distances were from the best expansion origins in previous research for the respective variables (Cenac, 2022b, 2023b), and the number of permutations was set at 9,999 as in von Cramon-Taubadel (2019); this number of permutations was also used when running linear regressions on distance matrices.

The regression test for distance matrices is an outreach of the Mantel correlation test (Diniz-Filho et al., 2013); because the Mantel correlation test is not as able to detect true associations compared to a correlation test (a correlation test for linear trends) (Legendre & Fortin, 2010), we may expect that regression tests on distance matrices in Main Analysis 2 would be weak at detecting relationships.

*Main Analysis 3* involved testing whether the genetic distance-based measure falls with the advance of geographical distance from Africa. This analysis used 50 populations. Due to positive **spatial autocorrelation** (see *Results and discussion*; *Glossary*), a correlation analysis was run using the Spatial Analysis in Macroecology (SAM) Version 4.0 software of Rangel et al. (2010) – https://www.ecoevol.ufg.br/sam/. In SAM Version 4.0, degrees of freedom were adjusted (using the Clifford et al. way ultimately) for spatial autocorrelation (Rangel et al., 2010).^4^ Whilst Betti (2014) used a different method to adjust analysis for spatial autocorrelation, inspiration was drawn from their analysis to make the adjustment in the current study. Betti (2014) used multidimensional scaling on interpopulation geographical distances which led to coordinates which were used when adjusting for spatial autocorrelation – this was so that the geographical distance between populations would reflect the smallest *travelling* distances rather than the shortest *physical* distance between populations. For instance (e.g., given Henn et al., 2012), the travelling distance from southern Africa to South America would be far larger than the physical distance between southern Africa and South America. Inspired by Betti (2014), multidimensional scaling was used (in R) on interpopulation geographical distances, resulting in coordinates for two axes; these coordinates were used in SAM Version 4.0 when adjusting degrees of freedom for spatial autocorrelation. There was a follow-up analysis of sorts. This involved reperforming analysis from Cenac (2023b) regarding the cranial shape distance-based measure for males, but in the absence of Oceanian populations. Therefore, whilst Cenac (2023b) used 28 populations, the present study used 21 (in the follow-up analysis). As before (Cenac, 2023b), RMET 5.0 (Relethford & Blangero, 1990) was utilised to determine interpopulation cranial shape distances, and Spearman correlation coefficients were used to represent relationships between geographical distance and cranial shape distance.

*Main Analysis 4* was about predicting distance from Africa from two variables (the genetic distance-based measure and genetic diversity). Once more, 50 populations were used in linear regression. Forty-nine of the populations were HGDP-CEPH (Balloux et al., 2009; Friedlaender et al., 2008) and were therefore matched. The remaining population regarding diversity was the Melanesian population in the HGDP-CEPH (Balloux et al., 2009) and the remaining population regarding genetic distance was the Nasioi population (Friedlaender et al., 2008). Melanesians in HGDP-CEPH correspond to Nasioi (Friedlaender et al., 2008) and they were therefore matched. Additionally, the analysis was repeated (unplanned analysis) with the addition of mitochondrial diversity and X-chromosomal diversity. As part of the unplanned analysis, the ridge package (Cule et al., 2022) was used in R to run a ridge regression.

Positive spatial autocorrelation was tested for through the Chen (2016) spreadsheet (three-step method) which calculates a spatial Durbin Watson (*DW*) statistic. Bounds for the *DW* statistic are applicable to the spatial *DW* (Chen, 2016), and bounds (Savin and White, 1977) were indeed applied in the present study. In case of positive spatial autocorrelation, or if it was unclear if there was positive spatial autocorrelation, graphs generated in the Chen (2016) spreadsheet were consulted, and it was seen if excluding a population (or populations) from analysis would suggest that the positive spatial autocorrelation is absent.

A Bonferroni correction (for a discussion, see, e.g., VanderWeele & Mathur, 2019) was applied. Essentially, to have a family-wise error rate of .05, the alpha-level of .05 was partitioned between the four Main Analyses (with *p*-values generally being multiplied by four); i) for the further linear regression and six Mantel tests under Main Analysis 2, the *p*-values were multiplied by sixteen, ii) in Main Analysis 3, *p*-values were multiplied by eight, iii) in Main Analysis 4, *p*-values were also multiplied by eight. Outliers were defined as datapoints which, after being *z*-score-transformed, were at values in excess of |3.29| (e.g., Field, 2013).

### Model comparison

Like in earlier research (e.g., Manica et al., 2007), the Bayesian information criterion (BIC) was used to compare models. As previously (e.g., Betti et al., 2009; Manica et al., 2007), a model was defined as being as good as another model if they were within four BICs of each other. Indeed, model comparisons using the BIC can be used to indicate the origin of the global expansion (e.g., Manica et al., 2007). Essentially, locations across the world were utilised like each was where the expansion arose from (i.e., a different origin is featured in each model) (e.g., Betti et al., 2009; Manica et al., 2007). We find the model giving the lowest BIC and models within four BICs of this best model – the origin of the expansion is indicated to be someplace within the geographical area covered by i) the origin used in the best model and ii) the origins used in the models which produced BICs which were within four BICs of the best model (Manica et al., 2007). This procedure was followed i) in Main Analysis 1 to see whether the geographical area covers a smaller area with the inclusion of the additional cranial variable/variables, ii) in Main Analysis 3 in order to indicate the geographical area identified by the genetic distance-based measure, and iii) in Main Analysis 4 to find whether the geographical area is reduced with the inclusion of the genetic distance-based measure, and also in Main Analysis 4 to find the geographical area indicated by mitochondrial diversity. A formula (Masson, 2011) was used to calculate BICs.

In Main Analysis 1, the best model was found for predicting geographical distance from cranial shape diversity, and it was seen if the addition of cranial sexual size dimorphism and the cranial shape distance-based measure (as predictors) improved the prediction of geographical distance. Similarly, in Main Analysis 4, the best (lowest) BIC was found regarding the model predicting geographical distance from autosomal diversity, and it was seen whether the addition of the genetic distance-based measure (as a predictor) improved the prediction of geographical distance. In the unplanned part of Main Analysis 4, the best of the BICs was found for predicting geographical distance from various diversities together (autosomal, mitochondrial, and X-chromosomal), and it was tested whether prediction is improved by the addition of the genetic distance-based measure. The location which has the lowest BIC varies depending on the variables used to calculate the BICs (Cenac, 2022b). Therefore, the addition of variables could change the origin which gives the lowest BIC. Hence, to assess if the addition of variables improved model fit, BICs were also used to compare models regarding the origin which gave the lowest BIC for the model which had the additional variable(s).

As in Atkinson (2011), the model producing the lowest BIC was used when assessing if there was a significant relationship (between geographical distance and a biological variable). The BIC can be calculated from *R*^2^ (e.g., Masson, 2011); when linear regressions were used on distance matrices (Main Analysis 2), the model resulting in the greatest *R*^2^ was used to see whether there were significant relationships.

## Results and discussion

### Main Analysis 1

A linear regression was run for predicting distance from Africa from three cranial variables: shape diversity, sexual size dimorphism, and the shape distance-based measure. Using 26 populations, it was not clear if positive spatial autocorrelation was present, spatial *DW* = 1.53, which a graph (produced using the Chen, 2016, spreadsheet) suggested was attributable to the North Japan population (and possibly the Berg population and, to a lesser extent, perhaps the Zalavár population) in the Howells (e.g., 1996) data. Upon the exclusion of North Japan from analysis, the situation was still unclear, spatial *DW* = 1.53, with a graph suggesting this was possibly being driven by Berg and (possibly to a lesser level) Zalavár populations.^5^ Without Berg (and North Japan), positive spatial autocorrelation was clearly absent, spatial *DW* = 1.84. Collectively, the three cranial variables explained a significant amount of variation in distance from Africa, *F*(3, 20) = 11.65, *p* < .001, *R*^2^ = .64. In the regression (i.e., when considering the three cranial variables simultaneously), cranial sexual size dimorphism had a unique relationship with distance from Africa, *b* = 517,717.00 (*SE* = 181,175.50), *t*(20) = 2.86, *p* = .039 (Figure 3A); unique relationships with distance from Africa were not found for either cranial shape diversity, *b* = −13,566.40 (*SE* = 17,399.00), *t*(20) = −.78, *p* = 1.00 (Figure 3B), or the cranial shape distance-based measure, *b* = −12,470.20 (*SE* = 6,677.00), *t*(20) = - 1.87, *p* = .31 (Figure 3C). Therefore, amongst cranial variables, just cranial sexual size dimorphism appeared to have a unique portion of the expansion signal.

**Figure 3.**
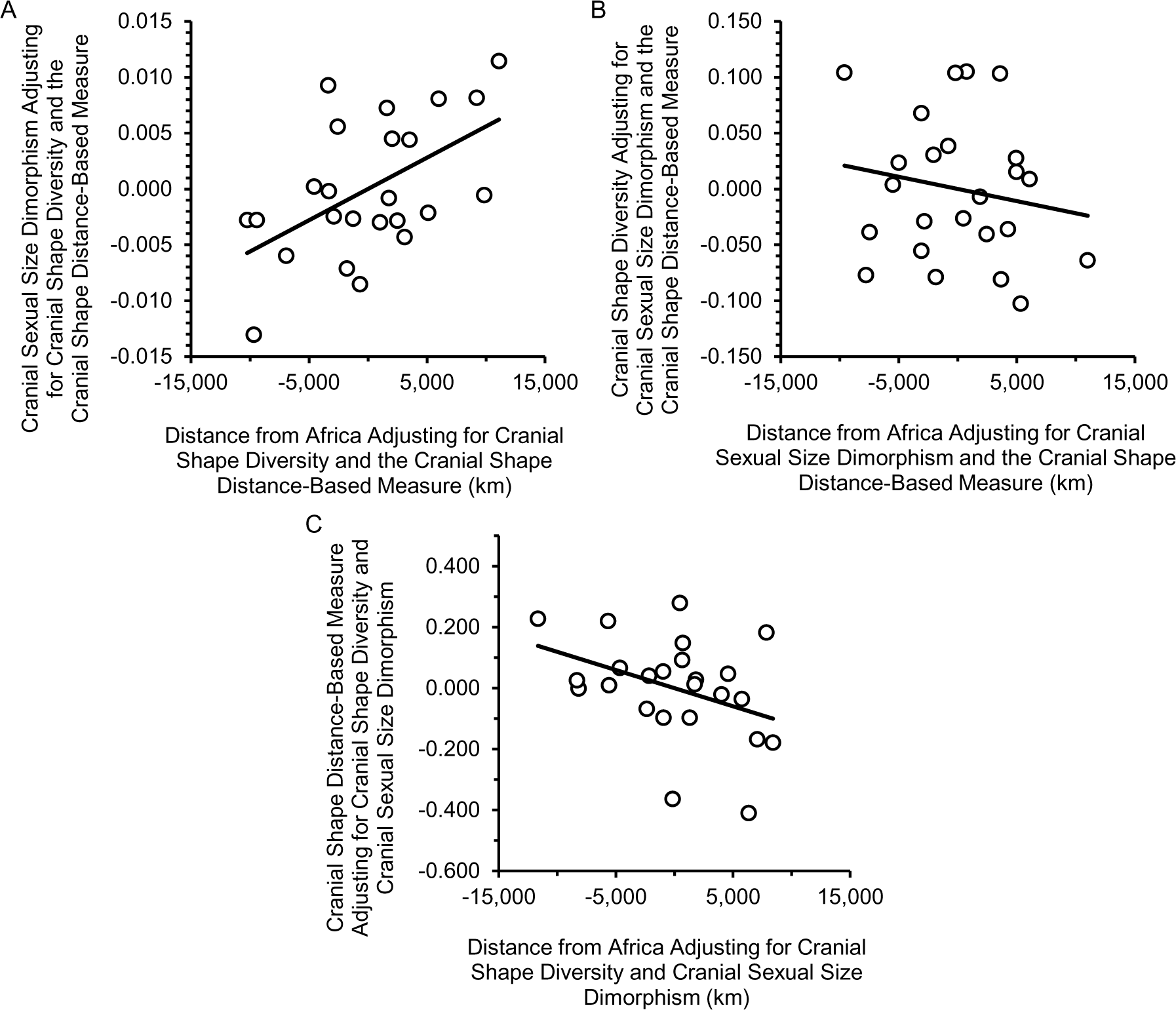
Partial Relationships Between Distance from Africa and Cranial Variables. *Note*. This figure shows partial relationships (i.e., controlling for other variables) between geographical distance from Africa and cranial sexual size dimorphism (3A), cranial shape diversity (3B), and the cranial shape distance-based measure (3C).

Cranial sexual size dimorphism is to do with size (Jantz & Ousley, 2020), so, with the other two cranial measures being shape-related (Cenac, 2023b; von Cramon-Taubadel & Lycett, 2008), there should have been an expectation for cranial sexual size dimorphism to be the variable which is most likely to have a unique part of the expansion signal.

Nonetheless, when using the location providing the lowest BIC for the relationship between cranial shape diversity and distance from Africa, BICs demonstrated that the model using the three cranial variables (*R*^2^ = .63) did not predict distance from Africa better than the model which only used cranial diversity (*R*^2^ = .47), *BIC*_Difference_ = 2.63. Similarly, when the location giving the lowest BIC when all three cranial variables was used, BICs showed that the model featuring all the cranial variables (*R*^2^ = .64) did not improve prediction of distance from Africa compared to the model which just featured cranial diversity (*R*^2^ = .47), *BIC*_Difference_ = 2.83. Therefore, despite cranial sexual size dimorphism having a unique part of the expansion signal, the expansion signal is indicated to not be recovered better with the inclusion of dimorphism and the distance-based measure. However, with only 24 populations being used when comparing BICs, it could be that comparisons were underpowered. Nevertheless, the signal of the expansion in the cranium (represented by the *R*^2^ of .64) still lags behind the signal which is present in genetic diversity (see Main Analysis 4).

Whilst inclusion of cranial sexual size dimorphism and the cranial shape distance-based measure did not seem to improve the recovery of the expansion signal, their inclusion still reduced the estimate for the geographical area which holds the origin of the expansion by 45.65% (Figure 2D).

This decreased area covered southern Africa, but went somewhat outside of that region too (e.g., Choudhury et al., 2021). Therefore, between positions of whether the south or the east of Africa was the origin of the expansion (Henn et al., 2018), the cranium favours the south over the east.^6^

The decrease in area is likely more attributable to cranial sexual size dimorphism than the cranial shape distance-based measure. This can be said for two reasons. Firstly, when considering the three cranial variables, only dimorphism exhibited a unique relationship with distance from Africa. Secondly, to infer which of the two variables was more responsible for the decrease in geographical area, BICs were used to estimate which area the origin is in (and not in) when using cranial shape diversity and i) cranial sexual size dimorphism (Figure 2B) or ii) the cranial shape distance-based measure (Figure 2C). Compared to the use of cranial shape diversity alone, the inclusion of cranial sexual size dimorphism reduced the area by 28.26% (Figure 2B), whilst the inclusion of the cranial shape distance-based measure decreased the area by only 10.87% (Figure 2C). These decreases appear to fall short of the near-50% decline when both variables were added (Figure 2D), suggesting that the addition of both variables together was particularly useful for reducing the area.

### Main Analysis 2

The ecodist package (Goslee & Urban, 2007) was used to run a linear regression, predicting pairwise (interpopulation) difference in geographical distance from Africa using four interpopulation distance (difference) variables: i) autosomal microsatellite heterozygosity, ii) cranial shape diversity, iii) cranial sexual size dimorphism, and iv) the cranial shape distance-based measure. In the model, difference in distance from Africa was uniquely related to difference in autosomal microsatellite heterozygosity (*p* < .001), but not to differences in either cranial shape diversity (*p* = 1.00), cranial sexual size dimorphism (*p* = 1.00), or the cranial shape distance-based measure (*p* = .74). This could suggest that the portions of the expansion signal in the cranial measures are not unique when considered alongside each other and autosomal microsatellite heterozygosity.

Is the portion of the signal in cranial shape diversity redundant specifically to the signal in autosomal diversity? Given that a unique part of the expansion signal is not present in cranial shape diversity when cranial shape diversity is considered alongside the other two cranial variables (Main Analysis 1), it could be that the portion of the signal in cranial shape diversity (which is not in autosomal diversity) was accounted for in cranial sexual size dimorphism and/or the cranial shape distance-based measure. Therefore, a further linear regression was run on matrices using ecodist (post-hoc) in the absence of the dimorphism and distance-based variables; a unique portion of the expansion signal was indicated for autosomal diversity (*p* = .003), but not for cranial shape diversity (*p* = 1.00). This seems congruent with the expansion signal in cranial shape diversity indeed being redundant to the signal in autosomal diversity.

However, analyses used only nine Howells populations; it could be that the expansion signal just happens to not appear present amongst the crania of those nine populations. Therefore, it was important to run Mantel tests (for relationships between geographical distance and biological distance) with a greater number of populations than the nine (of which the nine populations are a subset), and it was also important to run Mantel tests with the nine populations alone.

Results when using the nine populations alone will be addressed first; Mantel correlation tests were run on differences in geographical distance from Africa and i) differences in cranial shape diversity (*r*_Mantel_ = .35, *p* = 1.00) ii) differences in cranial sexual size dimorphism (*r*_Mantel_ = .15, *p* = 1.00), and iii) differences in the cranial shape distance-based measure (*r*_Mantel_ = .13, *p* = 1.00) – no significant relationships were found.^7^ Therefore, it should not be surprising for the linear regressions to not be indicative of unique parts of the expansion signal for any of the cranial variables. And so, it is not clear if the cranial measures have portions of the expansion signal which are distinct to the portion of the signal in autosomal microsatellite heterozygosity.

What about the Mantel tests that used more than the nine populations? Mantel tests were run using 24 populations with respect to cranial sexual size dimorphism (two populations were absent due to positive spatial autocorrelation in their presence in Cenac, 2022b),^8^ and 28 populations with respect to cranial shape diversity and the cranial shape distance-based measure. Regarding dimorphism, there was a positive and significant correlation, *r*_Mantel_ = .43, *p* = .006. For shape diversity, a positive, significant correlation was also observed, *r*_Mantel_ = .41, *p* = .002. However, a significant correlation was not evident with respect to the cranial shape distance-based measure, *r*_Mantel_ = .15, *p* = 1.00. Consequently, these results support a portion of the expansion signal being present in cranial sexual size dimorphism and cranial shape diversity; for those two variables, the nine populations perhaps just so happened to not capture the expansion signal. Nevertheless, results call into question whether the signal truly is there in the cranial shape distance-based measure.

Regarding the cranial shape distance-based measure, nonindependence was a concern in (Pearson) correlation analysis (Cenac, 2023b); assuming that the Mantel test handles nonindependence (e.g., Balloux et al., 2009; but see Guillot & Rousset, 2011), the nonsignificant result of the Mantel test could suggest that there actually is no expansion signal in the cranial shape distance-based measure. However, the Mantel test is less apt at finding actual relationships than a correlation analysis (linear) (Legendre & Fortin, 2010), with the Mantel test giving a smaller coefficient (Quilodrán et al., 2023).

### Main Analysis 3

Regarding the possible relationship between the genetic distance-based measure and geographical distance, the lowest BIC was found using an origin in Asia (Tel Aviv, Israel). Positive spatial autocorrelation was apparent, spatial *DW* = .99, and inspection of a graph (in the Chen, 2016, spreadsheet) suggested that the autocorrelation was prevalent across datapoints. Therefore, a correlation test was run using SAM Version 4.0, through which an adjustment to the degrees of freedom was applied (the degrees of freedom would have been 48 if not adjusted); as distance from Asia increased, there was not found to be a decline in the genetic distance-based measure, *r*(6.95) = - .80, *p* = .080 (Figure 4A).^9^ This may seem odd as the cranial shape distance-based measure seemed congruent with the expansion before (Cenac, 2023b), and the expansion signal is stronger at the genetic level than in the cranium (e.g., Roseman & Weaver, 2007), so we may have expected there to be a clear expansion signal in the genetic distance-based measure. Therefore, the absence of a signal calls into question whether the cranial shape distance-based measure really reflects the expansion.

**Figure 4.**
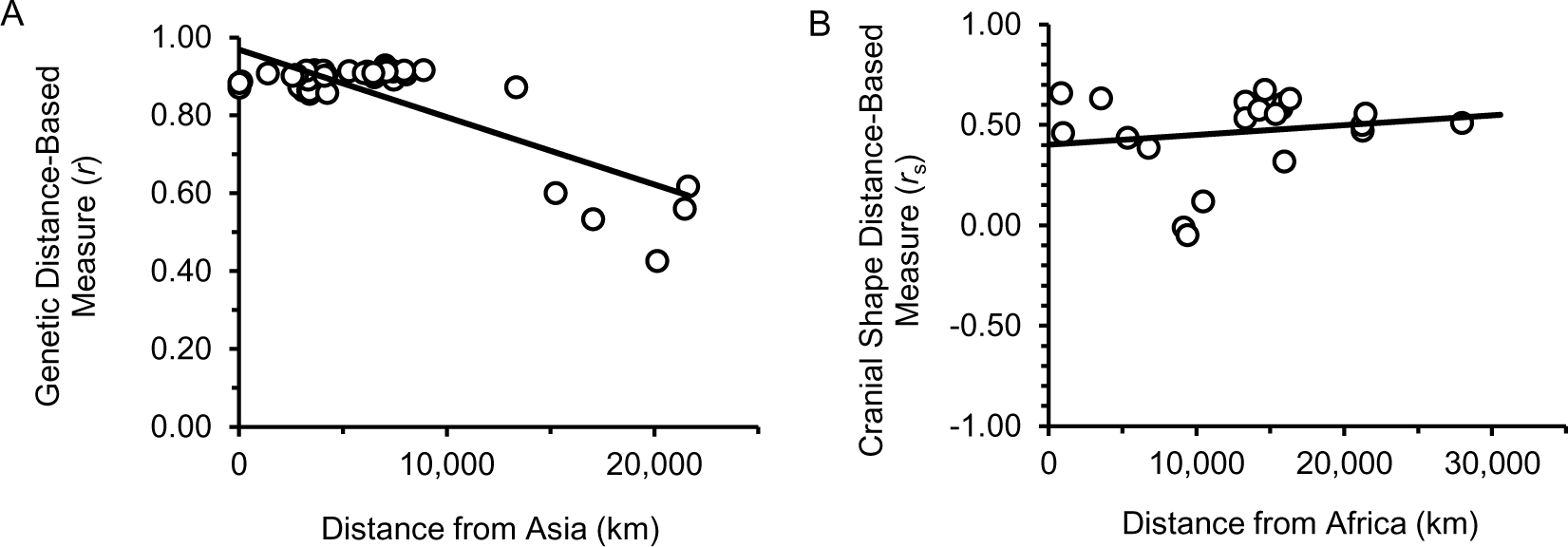
The Genetic Distance-Based and Cranial Shape Distance-Based Measures at Geographical Distances. *Note*. Figure 4A is with respect to the genetic distance-based measure whilst Figure 4B concerns the cranial shape distance-based measure. In Figure 4A, only American populations (López Herráez et al., 2009) had *y*-axis values under *r* = .80. Figure 4B is without Oceanian populations.

Even if the adjustment to the degrees of freedom was overly conservative, there would be some reason to doubt that a global expansion is reflected in the supposed relationship between the genetic distance-based measure and geographical distance from Asia. Variables which possess the signal of the global expansion exhibit a gradual change with distance from the origin (e.g., Betti et al., 2009; Prugnolle et al., 2005); the genetic distance-based measure does not indicate a gradual change, but roughly similar values except for the American populations who seem to be notably lower in the measure in comparison to the other populations (Figure 4).

From previous research (Cenac, 2023b; Howells, 1989; López Herráez et al., 2009; von Cramon-Taubadel & Lycett, 2008), one difference between the data used for the genetic distance-based measure (present study) and the cranial shape distance-based measure (in Cenac, 2023b) is that the cranial variable proportionally featured far more Oceanian populations – the genetic distance-based measure had one Oceanian population out of 50 populations (2.00%), whilst Cenac (2023b) used seven Oceanian populations out of a total of 28 population (25.00%). Therefore, it was decided to replicate analysis in Cenac (2023b) regarding whether the cranial shape distance-based measure is correlated with geographical distance from Africa, however, this time around, the analysis did not include Oceanian populations. Whilst the measure decreases with distance from Africa when Oceanian populations are featured (Cenac, 2023b), the present study found that, in their absence, there was no decrease (and indeed no significant relationship) between the measure and distance from Africa, *r*(19) = .16, *p* = 1.00 (Figure 4B). Yet, it was unclear whether positive spatial autocorrelation was at work, spatial *DW* = 1.25. Nonetheless, spatial autocorrelation promotes Type I errors, i.e., the wrong refutation of the null hypothesis (Deblauwe et al., 2012). Therefore, the failure to find a correlation between the cranial distance-based measure and distance from Africa without Oceanian populations (Figure 4B) would stand even if the positive spatial autocorrelation was corrected for. Therefore, results suggested that the decline found previously (Cenac, 2023b) was greatly attributable to the inclusion of Oceanian populations. By extension, it could be that the absence of an expansion signal in the genetic distance-based measure could be due to the sparse representation of Oceanians.

### Main Analysis 4

A linear regression was used for predicting distance from Africa from autosomal diversity and the genetic distance-based measure. When analysis featured the Bedouin HGDP-CEPH population (Balloux et al., 2009; Friedlaender et al., 2008), it was not clear if there was positive spatial autocorrelation, spatial *DW* = 1.52. A graph (using the Chen, 2016) spreadsheet suggested that this uncertainty arose from the Bedouin population, and, in their absence positive spatial autocorrelation was absent, spatial *DW* = 1.73; a linear regression suggested that a model using autosomal diversity and the genetic distance-based measure recovered the expansion signal, *F*(2, 46) = 173.50, *p* < .001, *R*^2^ = .88. Autosomal diversity had a unique relationship with distance from Africa, *b* = −86,603.00 (*SE* = 8,592.00), *t*(46) = −10.08, *p* < .001 (Figure 5A), as did the genetic distance-based measure, *b* = - 10,843.00 (*SE* = 3,704.00), *t*(46) = −2.93, *p* = .042 (Figure 5B). Regarding the genetic distance-based measure, the significant relationship with geographical distance in Main Analysis 4 appeared to be in contrast to Main Analysis 3, in which a significant relationship was not observed. The difference in results could seem attributable to autosomal diversity being adjusted for in Main Analysis 4 when gauging if there is a relationship between the genetic distance-based measure and geographical distance.

**Figure 5.**
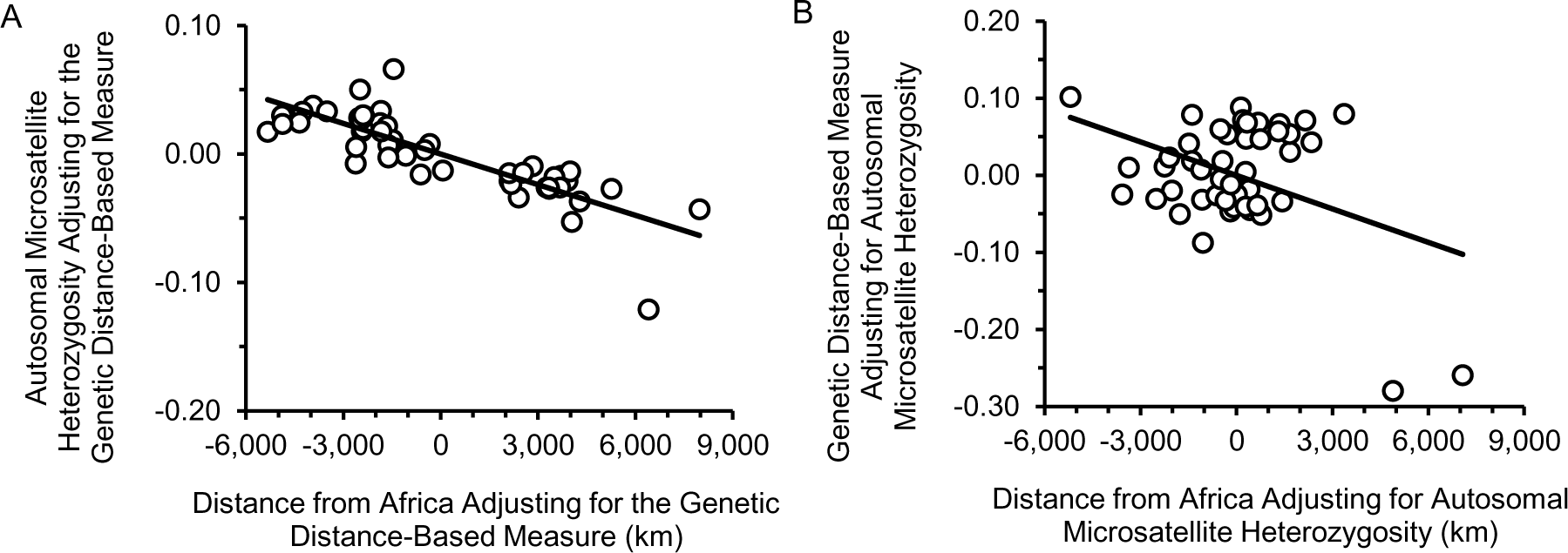
Partial Relationships Between Distance from Africa and Genetic Variables. *Note*. Figure 5 displays partial relationships (i.e., with another variable adjusted for) between geographical distance from Africa and autosomal microsatellite heterozygosity (5A) and the genetic distance-based measure (5B).

However, we can see that the relationship between the genetic distance-based measure and distance from Africa (controlling for autosomal diversity) is driven by two datapoints (Colombian and Mayan populations, Balloux et al., 2009, Friedlaender et al., 2008) being lower than the other populations, with the bulk of the other populations forming a cloud, and the datapoints within this cloud do not seem to exhibit a trend (Figure 5B). When a variable reflects the expansion, we expect a steady relationship between the variable and distance from the *origin* (because of the impact of bottlenecks in the expansion) (e.g., Ramachandran et al., 2005), not constant values followed by a sharp change (Figure 5B and, indeed, Figure 4A).

The addition of the genetic distance-based measure (compared to using autosomal diversity alone) may seem to improve recovery of the expansion signal, or not, depending on which origin is used. If we use the origin which gives the lowest BIC regarding the relationship between autosomal diversity and geographical distance, then the addition of the genetic distance-based measure does not improve our prediction of distance from Africa (*R*^2^ = .88) compared to genetic diversity alone (*R*^2^ = .86), *BIC*_Difference_ = 3.72. On the other hand, if we use the origin which gives the lowest BIC regarding the prediction of distance from Africa from both genetic variables, then model fit is better when both variables are used (*R*^2^ = .88) than when just diversity is used (*R*^2^ = .86), *BIC*_Difference_ = 4.49. When using both genetic variables, as opposed to genetic diversity alone, there was a 31.92% reduction in the size of the area which the origin is estimated to be within (Figure 6A and 6B). However, these model comparisons and the area comparison in Main Analysis 3 could be meaningless because of the possible absence of an expansion signal in the genetic distance-based measure (Main Analyses 3 and 4).

**Figure 6.**
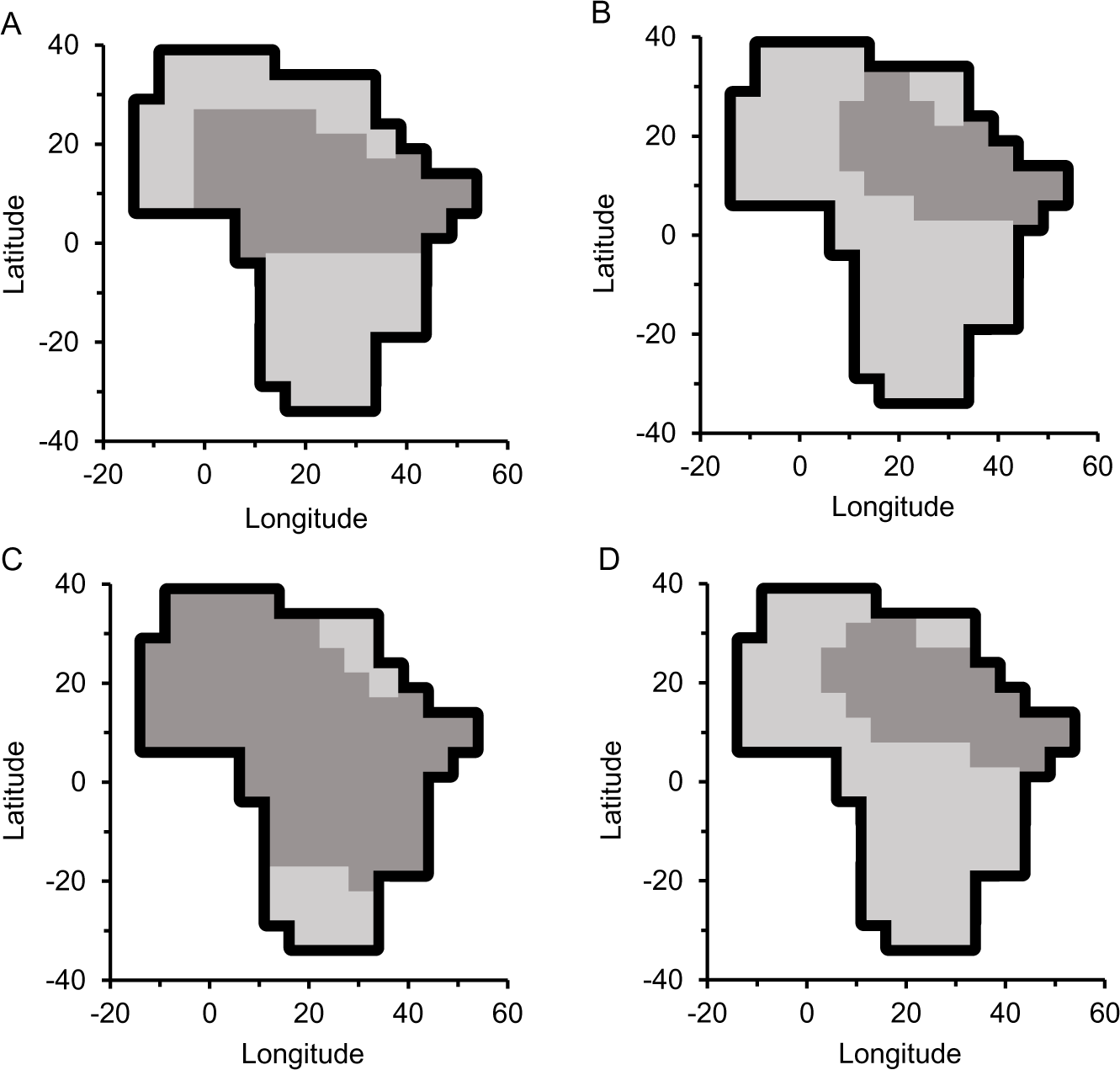
Genetics: Identification of Where the Expansion Originated? *Note*. As in Figure 2, the darker grey area signifies the area that the origin is in. Figure 6A was calculated using autosomal microsatellite heterozygosity, whilst Figure 6B was calculated using that diversity alongside the genetic (autosomal microsatellite) distance-based measure. Figure 6C used autosomal microsatellite heterozygosity, mitochondrial diversity, and X-chromosomal microsatellite heterozygosity, whereas Figure 6D used those diversities as well as the genetic distance-based measure.

Autosomal microsatellite heterozygosity is not the only genetic diversity which seems to have some of the expansion signal – for instance, portions of the signal seem to be present in X-chromosomal microsatellite heterozygosity and mitochondrial diversity too (e.g., Balloux et al., 2009); it was of interest to know if the genetic distance-based measure has a part of the signal which is distinct to the signal in not only autosomal diversity, but X-chromosomal and mitochondrial diversities too. When using a linear regression to predict distance from Africa from genetic diversities (autosomal, mitochondrial, and X-chromosomal) and the genetic distance based-measure, it was unclear whether positive spatial autocorrelation was present, spatial *DW* = 1.67. A graph produced in the Chen (2016) spreadsheet suggested that this was attributable to the Bedouin population; without that population, positive spatial autocorrelation appeared to be absent, spatial *DW* = 1.91. Therefore, analysis proceeded without Bedouin.

Multicollinearity was indicated regarding the relationship between i) autosomal diversity and ii) the other predictor variables in the model (*VIF* = 11.12). Similarly, multicollinearity was suggested when it came to the relationship between X-chromosomal diversity and the other predictors (*VIF* = 8.11). Multicollinearity was not suggested with respect to the relationship between the genetic distance-based measure and other predictors (*VIF* = 2.30) nor for the relationship between mitochondrial diversity and the remaining predictors (*VIF* = 1.44). Ridge regression can be used as a remedy for multicollinearity (e.g., Deng & Liu, 2022); through the ridge package, a ridge regression was applied in R (with lambda set to the automatic option, and the default scaling option chosen) (Cule et al., 2022) in place of a linear regression. As distance from Africa increased, significant decreases were observed with respect to three variables i) autosomal diversity, *b* = −16,831.55 (*SE* = 4,472.24), *t* = 3.76, *p* = .001, ii) X-chromosomal diversity, *b* = −12,586.02 (*SE* = 3,903.57), *t* = 3.22, *p* = .010, and iii) the genetic distance-based measure, *b* = −9,620.05 (*SE* = 2,458.40), *t* = 3.91, *p* = .001. As for mitochondrial diversity, analysis did not detect a significant variation with distance from Africa, *b* = 2,832.42 (*SE* = 2,006.84), *t* = 1.41, *p* = 1.00. Therefore, amongst the four variables, results suggested that there is a unique portion of the expansion signal in autosomal microsatellite heterozygosity and X-chromosomal microsatellite heterozygosity (thereby indicating that diversities can have different parts of the expansion signal), unlike mitochondrial diversity. As for the genetic distance-based measure, despite the ridge regression indicating that it has a unique part of the signal, there is clear reason to be skeptical about whether an expansion signal is present in this measure (e.g., Figures 4A and 5B).

Compared to when only the three genetic diversities were used (*R*^2^ = .87), inclusion of the genetic distance-based measure in analysis appeared to improve signal recovery (*R*^2^ = .90), *BIC*_Difference_ = 8.55, if geographical distance was from the location providing the lowest BIC when all four of the variables were used. Similarly, when geographical distance was from the place giving the lowest BIC when only the three diversities were used, the expansion signal seemed to be weaker without the genetic distance-based measure (*R*^2^ = .88) than with it (*R*^2^ = .90), *BIC*_Difference_ = 6.35. In contrast to when the three diversities were used, the addition of the genetic distance-based measure led to a 60.00% decrease in the area that the origin is estimated to be amongst (Figure 6C and 6D). However, yet again, there should be doubt over whether the genetic distance-based measure reflects the expansion (e.g., Main Analysis 3), which brings results into question.

Interestingly, the three genetic diversities do not collectively indicate that the origin is in any particular African region (Figure 6C). Different variables can deliver different perspectives of where the expansion originated (Manica et al., 2007). The estimate from genetic diversities is made of different variables (Figure 6C). X-chromosomal diversity and mitochondrial diversity have appeared (Cenac, 2022b) to give similar estimates to each other for the area which houses the origin. The area found with mitochondrial diversity in the present study (Figure 7) is similar to the area found previously (i.e., in Cenac, 2022b). Whilst there is some overal (to an extent), there are notable differences between the areas found with X-chromosomal or mitochondrial diversities (Cenac, 2022b; Figure 7) and the area found with autosomal diversity in the current study (Figure 6A). It could be that those differences resulted in a larger area being found when the three diversities were considered together than when only autosomal diversity was considered (Figure 6C and 6A). However, given previous preprints (Cenac, 2022b, 2023a), two points should be considered regarding autosomal diversity. The first point is linearity. It should be noted that analysis in the present study did not look for nonlinear trends. For autosomal microsatellite heterozygosity, a nonlinear (quadratic) trend encapsulates the relationship with distance from Africa to a greater degree than a linear trend does (Cenac, 2023a). Moreover, when it comes to defining the area that the origin may exist, an analysis allowing quadratic trends results in a geographical area containing the origin of the expansion (Cenac, 2023a) which seems similar to the areas (in Cenac, 2022b and Figure 7) for X-chromosomal and mitochondrial diversity. As for the second point, when using 106 populations of sub-Saharan Africans, the area holding the origin rests exclusively inside southern Africa (Cenac, 2022b).

**Figure 7.**
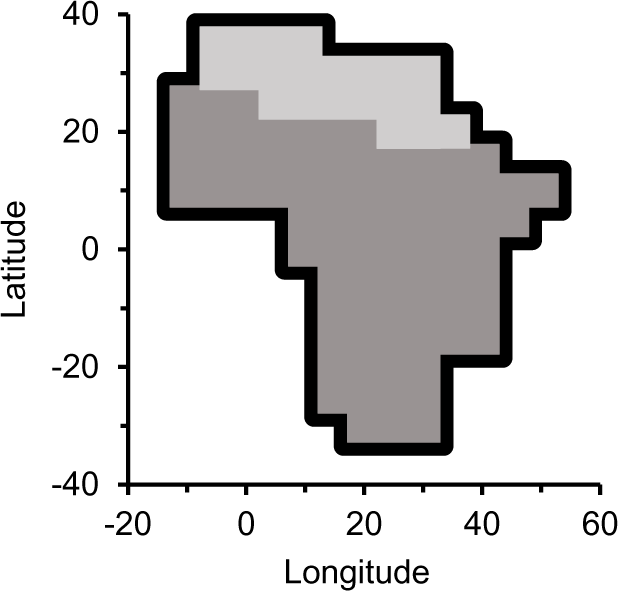
Identifying the Origin Through Mitochondrial Diversity. *Note*. The populations which were used were the ones in Main Analysis 4 when Bedouin were absent, i.e., 49 populations were used.

Even with shape diversity, sexual size dimorphism, and the shape distance-based variable being considered together, the expansion signal in the cranium does not appear to come close to matching the signal in genetics (Main Analyses 1 and 4). Because of this, the origin identified through genetics *ought* to be given more weight than the origin identified through the cranium. However, there were issues with identifying the origin through genetics in the present study (Figure 6), as reviewed above. Namely i) there is uncertainty over whether the genetic distance-based measure has part of the expansion signal, and ii) the origin identified in this study through autosomal microsatellite heterozygosity did not take nonlinearity into account, and it stands in contrast to the origin identified through this diversity beforehand using 106 sub-Saharan African populations (i.e., in Cenac, 2022b).^10^

## Conclusion

Overall, it was unclear if cranial sexual size dimorphism and biological distance-based measures are of any true value beyond diversity when identifying the origin of the expansion.^11^ Dimorphism and the cranial shape distance-based measure do not appear to improve recovery of the expansion signal (above cranial shape diversity), yet they do notably narrow the estimated range of locations in which the origin is situated (Figure 2), with this possibly being more attributable to dimorphism than the distance-based measure. Moreover, amongst cranial variables, only cranial sexual size dimorphism seems to have a unique portion of the expansion signal. If the cranial distance-based measure does have some of the expansion signal, then the combination of cranial variables gives its support to the origin being in the south more so than other African regions, however, support was not exclusive to the south (Main Analysis 1). Nevertheless, regarding biological distance-based measures, it is uncertain if they reflect the expansion (e.g., Main Analysis 3). The apparent presence of the expansion signal in biological distance-based measures may be contingent on Oceanian populations (Main Analysis 3). Therefore, it would be useful if future research supplements analysis of the genetic distance-based measure with more Oceanian populations. That research could make use of genetic distances regarding Oceanian populations which are available in Friedlaender et al. (2008) along with coordinates (and autosomal microsatellite heterozygosities) for those populations which were presented in Pemberton et al. (2013).

### Box 1

*Glossary*

- **Cranial shape distance-based measure**. Expansion from Africa gives a reason for why interpopulation (pairwise) genetic distance rises with the geographical distance between the populations (Ramachandran et al., 2005). The cranial distance between populations also increases (pairwise) with geographical distance (e.g., Betti et al., 2010; Hubbe et al., 2009). The cranial shape distance-based measure consisted of 28 correlations between interpopulation (pairwise) cranial shape distance (of males) and interpopulation (pairwise) geographical distance, with each correlation using distances from a particular population (e.g., Correlation 1 used pairwise distances between Population 1 and other populations, Correlation 2 used pairwise distances between Population 2 and other populations, etc.) – there were 28 populations, therefore 28 correlation coefficients (Cenac, 2023b). The idea is that the strength of correlations decreases as the population (which pairwise distance is from) gets farther from the origin, with this decrease happening because of the expansion being made of different routes (Cenac, 2023b).
- **Genetic distance-based measure**. Like the cranial shape distance-based measure, but using data from autosomal microsatellites rather than cranial shape data (see *Method*).
- **Spatial autocorrelation**. Spatial autocorrelation refers to, within a variable, datapoints being associated with datapoints that are geographically nearby – it can be negative (e.g., two datapoints which are geographically proximate are less similar than are two datapoints at random places), or positive (rather than being dissimilar, geographically proximate datapoints are similar) (e.g., Dubé & Legros, 2014). If errors/residuals in an analysis are spatially autocorrelated, then an assumption for regression analysis is not fulfilled – the assumption that errors are independent (e.g., Dubé & Legros, 2014).

## Footnotes

1 The procedure was that i) for each genetic/cranial variable, 99 locations in Africa were ranked (1 to 99) according to how good a fit they were for being the origin, ii) for each location, an average of the ranks was calculated across the variables, iii) these averaged ranks were reranked (1 to 99), thereby giving an impression of where the expansion originated across genetic and cranial variables (Cenac, 2023b).

2 Technically, a number of variables have been used simultaneously when exploring the origin of the expansion and carving out a distinctive area for the origin (e.g., Betti et al., 2009, 2013). For instance, when using cranial diversity, cranial diversity is calculated as the mean standardised diversity across a set of cranial measurements/variables (e.g., Betti et al., 2009; Manica et al., 2007). Nonetheless, in an analysis, cranial diversity is treated as *one* variable (e.g., Betti et al., 2009). And so, in the present study, *different variables* refers to when variables are treated as different variables in an analysis, e.g., cranial diversity and cranial sexual size dimorphism would be different variables.

3 This measure was used because Balloux et al. (2009) found that mitochondrial diversity (calculated as the average of pairwise disparities) declines with increased geographical distance from Africa (Balloux et al., 2009); mitochondrial diversity from Balloux et al. (2009) has been used to suggest global expansion from (somewhere within) an African geographical area alone (Cenac, 2022b).

4 In SAM Version 4.0, degrees of freedom in correlation analysis can be adjusted for the spatial autocorrelation *within* variables (Rangel et al., 2010). Dealing with the spatial autocorrelation within variables should go aways to handling the spatial autocorrelation amongst the residuals which were produced by the model that was used for predicting geographical distance from the genetic distance-based measure. SAM Version 4.0 gives two ways of adjusting the degrees of freedom – Dutilleul (1993) and Clifford et al. (1989) (Rangel et al., 2010). Both ways were applied, and the one leading to the most conservative decrease in the degrees of freedom (the Clifford et al. route) was selected in order to minimise the chance of Type I errors by the most.

5 The nearer populations are to one another, the more alike they are in their genetics (e.g., Ramachandran et al., 2005), i.e., spatial autocorrelation is evident regarding genetics (Creanza et al., 2015). Moreover, with geographical closeness also comes cranial similarity (Betti et al., 2010; Hubbe et al., 2009). Therefore, some level of positive spatial autocorrelation (amongst errors) in the present study should not be surprising. Indeed, when it comes to the relationship between cranial sexual size dimorphism (calculated from the Howells data) and geographical distance from Africa, we can see a positive spatial autocorrelation which seems to be caused by Ainu and North Japan populations (Cenac, 2022b), and these populations are comparatively close to each other, only being 98.17 km apart (Cenac, 2022a); this suggests/implies that positive spatial autocorrelation only became notable regarding dimorphism when populations were particularly close to each other. In Main Analysis 1, it was uncertain if positive spatial autocorrelation was at work, and, once again, this involved the North Japan population, but additionally Berg and maybe Zalavár populations. Berg and Zalavár are just 296.63 km apart (Cenac, 2022b).

6 When using all 26 populations, and without North Japan, the lowest two BICs were in Australia and Tasmania locations (from von Cramon-Taubadel & Lycett, 2008). It should be noted that the lowest BIC can be found away from the origin of the expansion – for mitochondrial diversity, the lowest is in North America, yet that North American location produces a positive (rather than a negative) relationship between mitochondrial diversity and geographical distance, which indicates the importance of taking into account the direction of relationships when identifying the origin of the expansion (Cenac, 2022b). Inconsistent with being the origin of the expansion, Oceania has i) positive correlation coefficients regarding cranial shape diversity, ii) negative correlation coefficients with respect to cranial sexual size dimorphism (Cenac, 2022b), and iii) positive correlation coefficients when it comes to the cranial shape distance-based measure (Cenac, 2023b). Therefore, those lowest BICs in Oceania were discarded.

7 Mantel tests have previously been used to verify relationships between distance from Africa and genetic diversities, in case of datapoints being nonindependent with respect to the relationships between distance from Africa and genetic diversities (Balloux et al., 2009).

8 The Mantel test suffers from an increased Type I error rate when spatial autocorrelation is present in both variables under analysis in the Mantel test (Guillot & Rousset, 2011). In research which tested whether cranial sexual size dimorphism varies with geographical distance from Africa, Ainu and North Japan were the Howells populations which seemed to be driving the positive spatial autocorrelation of residuals (Cenac, 2022b); these two populations are only at an estimated 98.17 km apart (Cenac, 2022a). Therefore, positive spatial autocorrelation (expressed at very short geographical distance) seems to be present in the cranial sexual size dimorphism variable itself. In the present study, the Mantel test which featured differences in cranial sexual size dimorphism also featured differences in geographical distance from Africa – it should not be surprising if this variable exhibits some level of spatial autocorrelation. Therefore, rather than running the Mantel test on 26 Howells data populations, only 24 were used. However, given the geographical distance between Ainu and North Japan, and that these populations seemed to be leading to positive spatial autocorrelation (Cenac, 2022a, 2022b), it may very well have been sufficient to not have included only one of these populations (either one) to eliminate spatial autocorrelation in previous research (i.e., Cenac, 2022a, 2022b) and when running the Mantel test in the present study.

9 Spatial autocorrelation can be dealt with in SAM Version 4.0 through spatial eigenvector mapping (Rangel et al., 2010). That method was used in SAM Version 4.0 to try to find eigenvectors that would neutralise the positive spatial autocorrelation in residuals by using a model for predicting distance from Asia from the genetic distance-based measure and six eigenvectors. However, it was not clear if residuals from this model were positively spatially autocorrelated, spatial *DW* = 1.65.

10 The relationship between geographical distance from Africa and mitochondrial diversity would seem to be represented better by a nonlinear (quadratic) relationship than a linear relationship (Cenac, 2023a). However, mitochondrial diversity has a climatic signal (Balloux et al., 2009); when climate is adjusted for, the nonlinear relationship with distance from Africa is no longer supported (Cenac, 2023a). Compared to analysis which only considers linear relationships, the consideration of nonlinear relationships delivers a smaller estimate for the geographical area which the origin is within (Cenac, 2023a), however, the geographical areas from linear and quadratic models (Cenac, 2022b, 2023a) are still roughly similar. This suggests that, whilst the current study did not adjust mitochondrial diversity for climate, adjusting for climate would likely not have had a notable effect on results.

11 Of course, cranial sexual size dimorphism (Cenac, 2022b), and possibly biological distance-based measures (Cenac, 2023b; present study), are not the only variables aside from diversity to have the expansion signal – the signal is observable in linkage disequilibrium (Jakobsson et al., 2008). However, linkage disequilibrium reflects genetic diversity (e.g., Rosenberg & Kang, 2015) at a level to which (Conrad et al., 2006) it seems doubtful if linkage disequilibrium would have a tangible part of the expansion signal which is not in genetic diversity.

